# The identification of potent and selective antibodies for Serine/threonine-protein kinase TBK1, for use in immunoblot, immunofluorescence and immunoprecipitation

**DOI:** 10.1101/2022.06.03.494699

**Authors:** Walaa Alshafie, Maryam Fotouhi, Irina Shlaifer, Aled M. Edwards, Thomas M Durcan, Peter S. McPherson, Carl Laflamme

**Author notes:** Authors contributed equally and are listed alphabetically (Walaa Alshafie, Maryam Fotouhi).

## Abstract

TBK1 is a serine-threonine kinase that has been linked to a number of diseases, including amyotrophic lateral sclerosis and frontotemporal dementia. Reproducible research on TBK1 has been hampered by the lack of well characterized antibodies. In this study, we characterized 11 commercial antibodies for immunoblot, immunofluorescence and immunoprecipitation, using a knock-out cell line as the control. For each application, we identified several potent and selective antibodies that will facilitate studies on TBK1.

## INTRODUCTION

The lack of robust characterization of research antibodies contributes to the reproducibility crisis [1]. Rather than generate new antibodies, given that there are more than five million antibodies on the commercial market (CiteAb.com), we elected to test the hypothesis that among them will be some high performing antibodies [2].

TBK1 regulates autophagy though phosphorylation of Optineurin [3] and the C9ORF72/SMCR8 complex [4]. Of note, both Optineurin [5] and the C9ORF72/SMCR8 complex [6, 7] are directly linked to amyotrophic lateral sclerosis and frontotemporal dementia. Moreover, TBK1 also phosphorylates LC3C, GABARAP-L2 [8] and AKT1 [9] promoting autophagy.

The endogenous localization of TBK1 at basal state and during autophagy remains yet to be identified. Moreover, TBK1 protein interactomes have been done with overexpression system, with the exception of one study [10]. TBK1 antibodies are key to address these unknowns.

To explore the availability of high-quality antibodies, we used an antibody characterization strategy [11] based on knockout (KO) cells to perform head-to-head comparisons of commercial antibodies in immunoblot (Western blot), immunoprecipitation and immunofluorescence applications. Here, we apply this approach to the TBK1 protein and identified specific antibodies for all tested applications, enabling biochemical and cellular assessment of TBK1.

## Materials and methods

### Antibodies

All TBK1 antibodies are listed in Table 1. Peroxidase-conjugated goat anti-mouse and anti-rabbit antibodies are from Thermo Fisher Scientific (cat. number 65-6120 and 62-6520). Alexa-555-conjugated goat anti-mouse and anti-rabbit secondary antibodies are from Thermo Fisher Scientific (cat. number A21424 and A21429).

### CRISPR/Cas9 genome editing

Cell lines used are listed in Table 2. U2OS *TBK1* KO clone was generated with low passage cells using an open-access protocol available on Zenodo.org: https://zenodo.org/record/3738361#.YIyeDu2SlaR. Two guide RNAs were used to introduce a STOP codon in the *TBK1* gene (sequence guide 1: UUUGAACAUCCACUGGACGA, sequence guide 2: CAAAUUAUUUGCUAUUGAAG).

**Table 2:** Summary of the Serine/threonine-protein kinase TBK1 antibodies tested.

| **Company** | **Catalogue number** | **Lot number** | **RRID**<br>**(Antibody Registry)** | **Clonality** | **Clone ID** | **Host** | **Concentration**<br>**(µg/µl)** | **Vendor's recommended applications** |
| --- | --- | --- | --- | --- | --- | --- | --- | --- |
| Bio-Techne | NB100-56705 | B-1 | AB_838420 | monoclonal | 108A429 | mouse | 1.00 | Wb, IF |
| Proteintech | 28397-1-AP | 00076443 | AB_2881132 | polyclonal | - | rabbit | 0.43 | Wb |
| Proteintech | 67211-1-Ig | 10013180 | AB_2882504 | monoclonal | 2D7B1 | mouse | 1.00 | Wb, IF |
| Thermo | PA5-17478 | VL3152289A | AB_10981817 | polyclonal | - | rabbit | not provided | Wb, IF, IP |
| Thermo | 703154 | 2274494 | AB_2848223 | recombinant-mono | JM42-11 | rabbit | 0.50 | Wb, IF |
| Abcam | ab12116 | GR3334526-1 | AB_298856 | monoclonal | 108A429 | mouse | 1.00 | Wb |
| Abcam | ab40676 | GR3275777-2 | AB_776632 | recombinant-mono | EP611Y | rabbit | 1.48 | Wb, IF |
| Abcam | ab109735 | GR3263881-3 | AB_10863562 | recombinant-mono | EPR2867(2)-19 | rabbit | 0.50 | Wb, IP |
| GeneTex | GTX113057 | 43481 | AB_11174793 | polyclonal | - | rabbit | 0.70 | Wb, IF |
| Cell Signaling Technology | 38066 | 1 | AB_2827657 | recombinant-mono | E8I3G | rabbit | not provided | Wb, IP, IF |
| Cell Signaling Technology | 3504 | - | AB_2255663 | recombinant-mono | D1B4 | rabbit | not provided | Wb, IP |

Wb= Western blot; IF= immunofluorescence; IP=immunoprecipitation

### Cell culture

Cells were cultured in DMEM high-glucose (GE Healthcare cat. number SH30081.01) containing 10% fetal bovine serum (Wisent, cat. number 080450), 2 mM L-glutamate (Wisent cat. number 609065, 100 IU penicillin and 100 μg/ml streptomycin (Wisent cat. number 450201).

### Antibody screening by immunoblot

Immunoblots were performed as described in our standard operating procedure [15]. Lysates were sonicated briefly and incubated 30 min on ice. Lysates were spun at ~110,000xg for 15 min at 4°C and equal protein aliquots of the supernatants were analyzed by SDS-PAGE and immunoblot. BLUelf prestained protein ladder from GeneDireX (cat. number PM008-0500) was used.

Immunoblots were performed with large 5-16% gradient polyacrylamide gels and transferred on nitrocellulose membranes. Proteins on the blots were visualized with Ponceau staining which is scanned to show together with individual immunoblot. Blots were blocked with 5% milk for 1 hr, and antibodies were incubated O/N at 4°C with 5% bovine serum albumin in TBS with 0,1% Tween 20 (TBST). Following three washes with TBST, the peroxidase conjugated secondary antibody was incubated at a dilution of ~0.2 μg/ml in TBST with 5% milk for 1 hr at room temperature followed by three washes with TBST. Membranes are incubated with ECL from Pierce (cat. number 32106) prior to detection with HyBlot CL autoradiography films from Denville (cat. number 1159T41).

### Antibody screening by immunoprecipitation

Immunoprecipitation was performed as described [16]. Antibody-bead conjugates were prepared by adding 1.0 μg of antibody to 500 ul of PBS with 0,01% triton X-100 in a microcentrifuge tube, together with 30μl of protein A- (for rabbit antibodies) or protein G- (for mouse antibodies) Sepharose beads. Tubes were rocked O/N at 4°C followed by several washes to remove unbound antibodies.

U2OS WT were collected in HEPES buffer (20 mM HEPES, 100 mM sodium chloride, 1 mM EDTA, 1% Triton X-100, pH 7.4) supplemented with protease inhibitor. Lysates are rocked 30 min at 4°C and spun at 110,000xg for 15 min at 4°C. One ml aliquots at 1.0 mg/ml of lysate were incubated with an antibody-bead conjugate for ~2 hrs at 4°C. Following centrifugation, the unbound fractions were collected, and beads were subsequently washed three times with 1.0 ml of HEPES lysis buffer and processed for SDS-PAGE and immunoblot on a 5-16% acrylamide gel.

### Antibody screening by immunofluorescence

Immunofluorescence was performed as described [17]. U2OS WT and *TBK1* KO were labelled with a green and a deep red fluorescence dye, respectively. The fluorescent dyes used are from Thermo Fisher Scientific (cat. number C2925 and C34565). WT and KO cells were plated on glass coverslips as a mosaic and incubated for 24 hrs in a cell culture incubator. Cells were fixed in 4% PFA (in PBS) for 15 min at room temperature and then washed 3 times with PBS. Cells were permeabilized in PBS with 0,1% Triton X-100 for 10 min at room temperature and blocked with PBS with 5% BSA, 5% goat serum and 0.01% Triton X-100 for 30 min at room temperature. Cells were incubated with IF buffer (PBS, 5% BSA, 0,01% Triton X-100) containing the primary TBK1 antibodies O/N at 4°C. Cells were then washed 3 × 10 min with IF buffer and incubated with corresponding Alexa Fluor 555-conjugated secondary antibodies in IF buffer at a dilution of 1.0 μg/ml for 1 hr at room temperature with DAPI. Cells were washed 3 × 10 min with IF buffer and once with PBS. Coverslips were mounted on a microscopic slide using fluorescence mounting media (DAKO).

Imaging was performed using a Zeiss LSM 880 laser scanning confocal microscope equipped with a Plan-Apo 40x oil objective (NA = 1.40). Analysis was done using the Zen navigation software (Zeiss). All cell images represent a single focal plane. Figures were assembled with Adobe Illustrator.

## RESULTS AND DISCUSSION

To identify a cell line that expresses adequate levels of TBK1 protein to provide sufficient signal to noise, we examined public proteomics databases [12, 13]. U2OS was identified as a suitable cell line and thus U2OS was modified with CRISPR/Cas9 to knockout the corresponding *TBK1* gene (Table 1) [14].

**Table 1.**
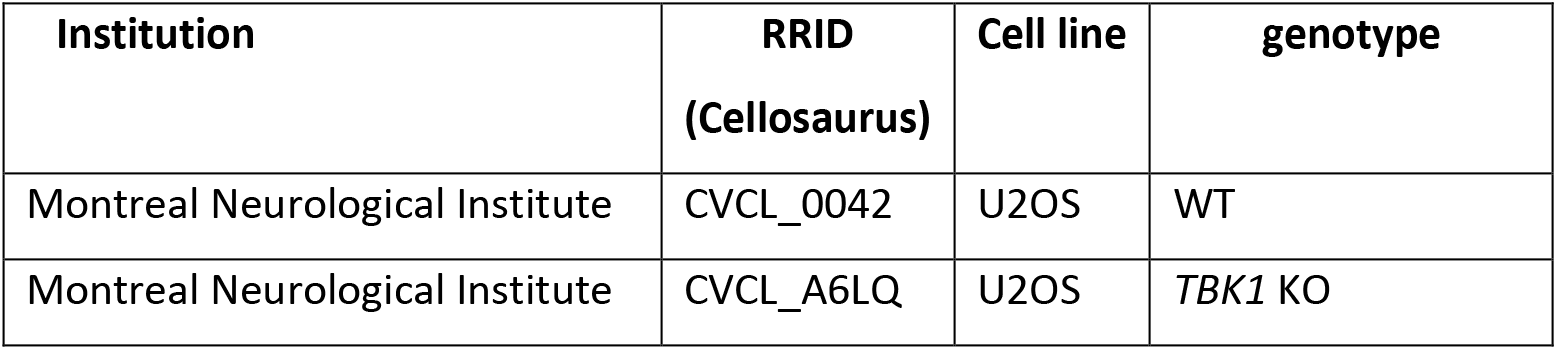
Summary of the cell lines used.

Extracts from wild-type and *TBK1* KO cells were prepared and used to probe the antibodies (Table 2) by immunoblot (Western blot) and immunoprecipitation, and the profile of each of the antibodies in immunoblot are shown in Figure 1 and in immunoprecipitation in Figure 2.

**Figure 1:**
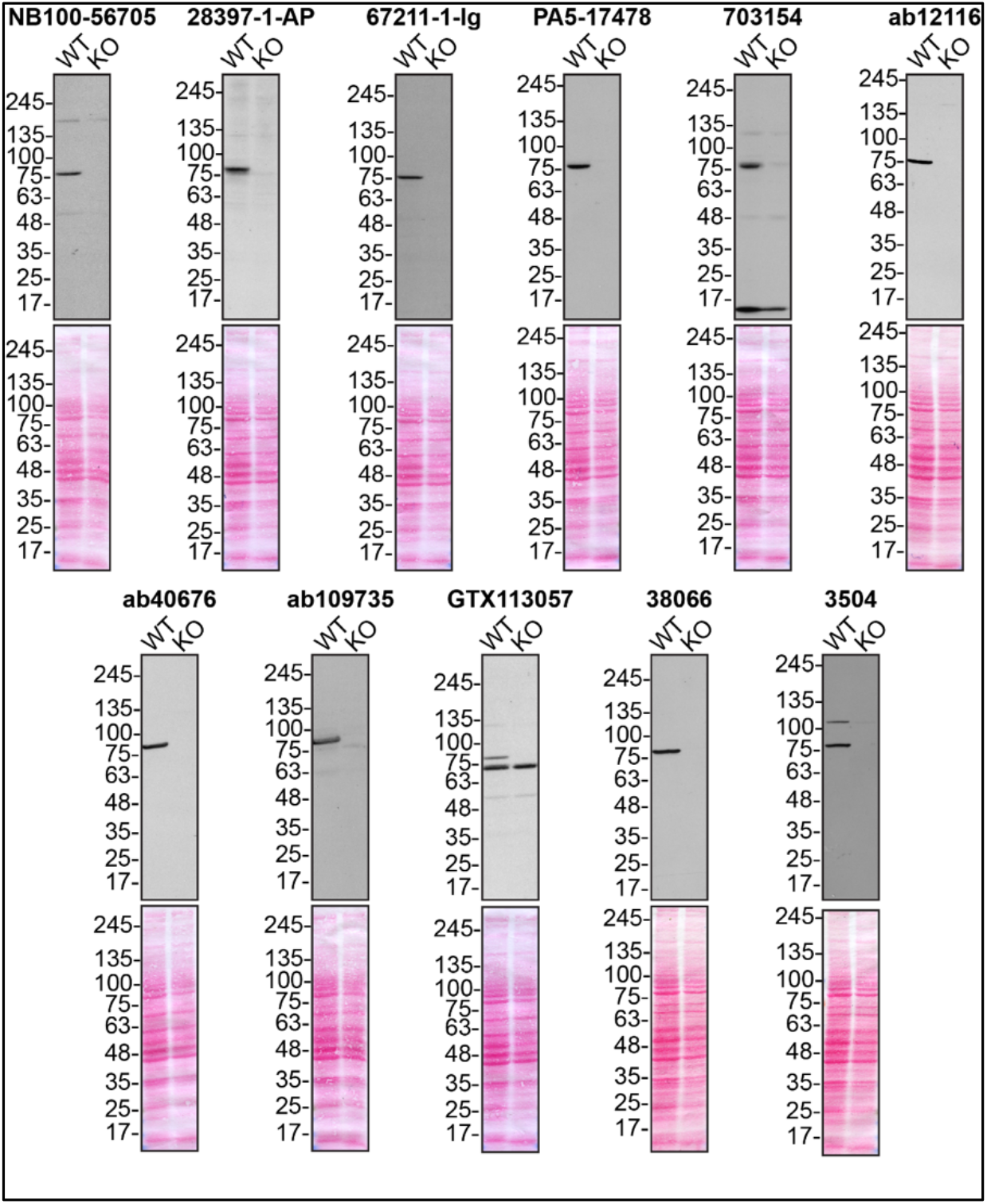
TBK1 antibody screening by immunoblot. Lysates of U2OS (WT and *TBK1* KO) were prepared and 50 μg of protein were processed for immunoblot with the indicated TBK1 antibodies. The Ponceau stained transfers of each blot are shown. Antibody dilution used was 1/5000 for all tested antibodies. Predicted band size: ~83 kDa.

**Figure 2:**
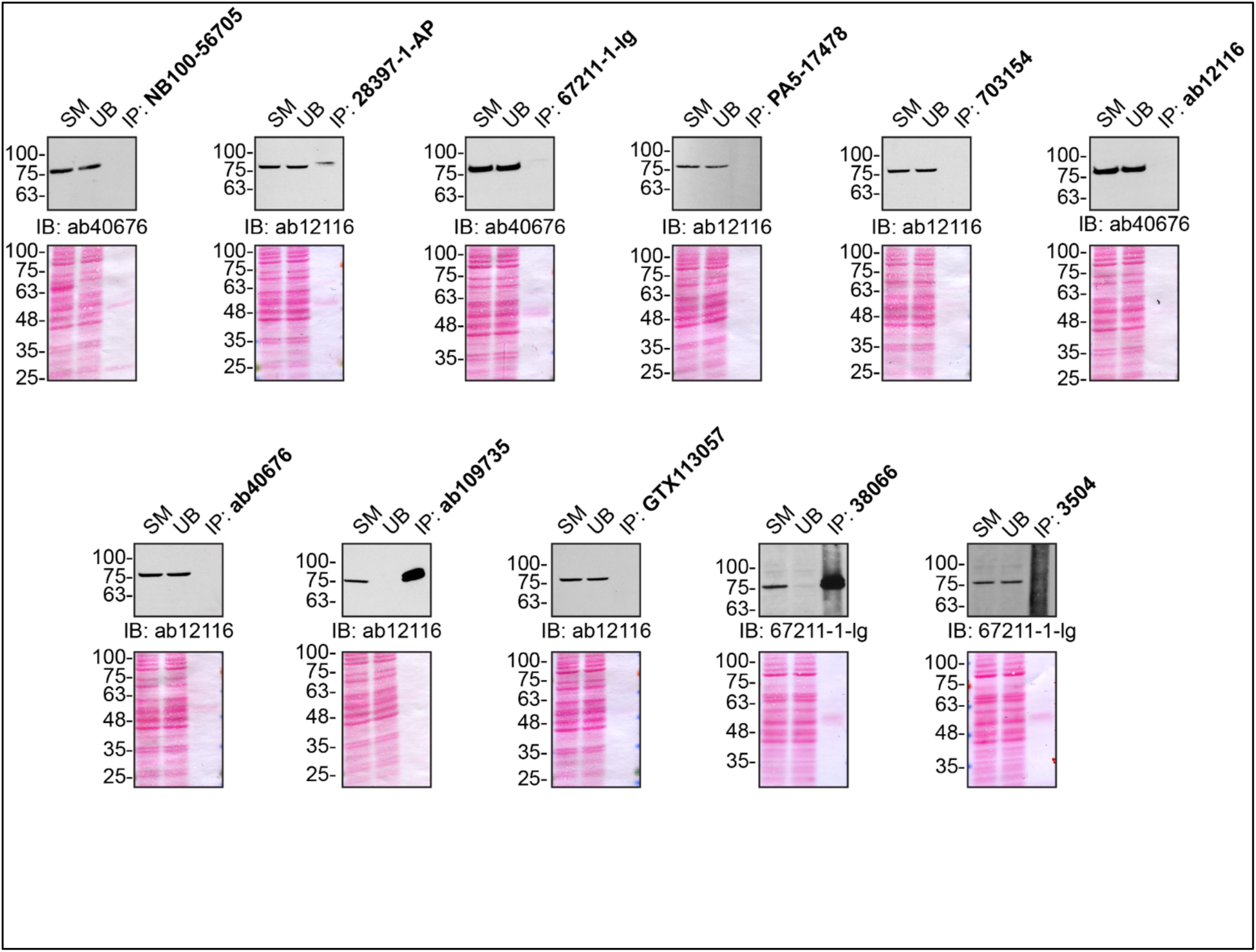
TBK1 antibody screening by immunoprecipitation. U2OS lysates were prepared and IP was performed using 1.0 μg of the indicated TBK1 antibodies precoupled to either protein G or protein A Sepharose beads. Samples were washed and processed for immunoblot with the indicated TBK1 antibody. For immunoblot, ab40676, ab12116 and 67211-1-Ig were used. The Ponceau stained transfers of each blot are shown. SM=10% starting material; UB=10% unbound fraction; IP=immunoprecipitate.

**Figure 3:**
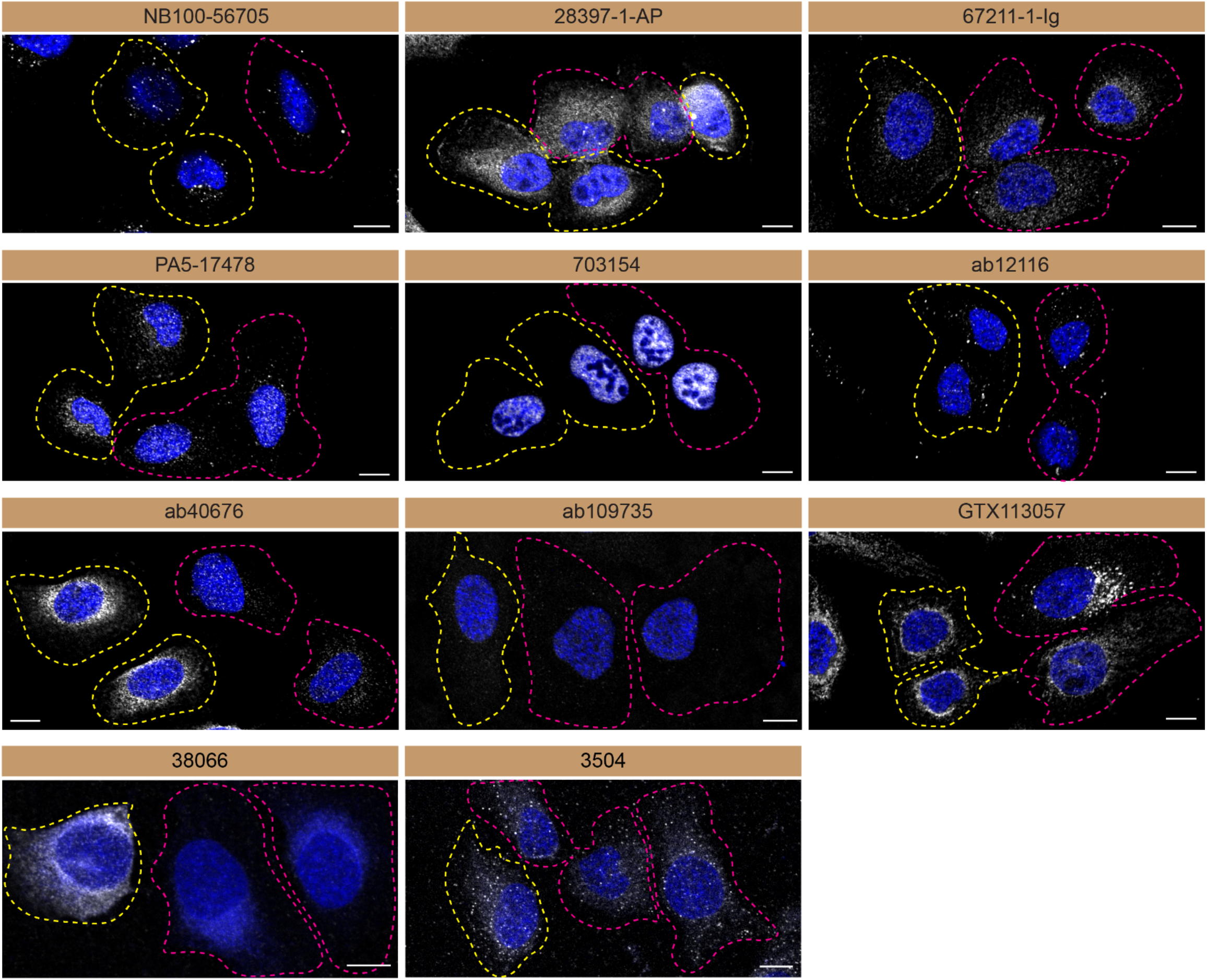
Serine/threonine-protein kinase TBK1 antibody screening by immunoblot. U2OS WT and *TBK1* KO cells were labelled with a green or a far-red fluorescent dye, respectively. WT and KO cells were mixed and plated to a 1:1 ratio on coverslips. Cells were stained with the indicated TBK1 antibodies and with the corresponding Alexa-fluor 555 coupled secondary antibody including DAPI. Acquisition of the blue (nucleus-DAPI), green (WT), red (antibody staining) and far-red (KO) channels was performed. Representative images of the merged blue and red (grayscale) channels are shown. WT and KO cells are outlined with yellow and magenta dashed line, respectively. Schematic representation of the mosaic strategy used is shown on the bottom-right panel. Antibody dilution used: NB100-56705 at 1/1000; 28397-1-AP at 1/500; 67211-1-Ig at 1/1000; PA5-17478 at 1/1000; 703154 at 1/500; ab12116 at 1/1000; ab40676 at 1/1500; ab109735 at 1/500; GTX113057 at 1/700, 38066 at 1/500, 3504 at 1/500. Bars = 10 μm.

For immunofluorescence, antibodies were screened using a mosaic strategy developed elsewhere [11]. Plating WT and KO cells together and imaging both cell type in the same field of view reduces imaging and analysis biases.

In conclusion, we have screened most TBK1 commercial antibodies by immunoblot, immunoprecipitation and immunofluorescence. The data provided can be used as a guide to purchase the most appropriate antibody for a researcher’s needs, and to support more reproducible research.

